# Application of a Novel Machine Learning Method to Big Data Infers a Relationship Between Asthma and the Development of Neoplasia

**DOI:** 10.1101/439117

**Authors:** Abbas Shojaee, Jose L. Gomez, Xiaochen Wang, Naftali Kaminski, Jonathan M. Siner, Seyedtaghi Takyar, Hongyu Zhao, Geoffrey Chupp

**Affiliations:** Section of Pulmonary, Critical Care and Sleep Medicine, Department of Internal Medicine, Yale University School of Medicine, New Haven, Connecticut, USA; Center for Precision Pulmonary Medicine (P2MED), Yale University School of Medicine, New Haven, Connecticut, USA; Department of Biostatistics, School of Public Health, Yale University, New Haven, Connecticut, USA

**Author notes:** Author for Correspondence: Dr. Geoffrey Chupp, Section of Pulmonary, Critical Care and Sleep Medicine, Department of Internal Medicine, Yale University School of Medicine, New Haven, Connecticut 06510. Authors Contribution: Abbas Shojaee designed the study, developed the methodology, performed analysis and wrote the manuscript, Geoffrey Chupp supervised the research and together with Naftali Kaminski and Jose Gomez verified the results and critically contributed to writing the manuscript, Seyedtaghi Takyar conceived the idea of the asthma-neoplasms relationship and contributed to the writing. Hongyu Zhao and Xiaochen Wang contributed to verifying methods and writing.

**Keywords:** Asthma, Neoplasm, Endocrine, Glandular Neoplasia, Etiology, Chronic Obstructive Pulmonary Disease, Malignancy

## Abstract

**Background:** A relationship between asthma and the risk of having cancer has been identified in several studies. However, these studies have used different methodologies, been primarily cross-sectional in nature, and the results have been contradictory. Population-level analyses are required to determine if a relationship truly exists.

**Methods:** We developed a novel machine learning tool to infer associations, Causal Inference using the Composition of Transactions (CICT). Two all payers claim datasets of over two hundred million hospitalization encounters from the US-based Healthcare Cost and Utilization Project (HCUP) were used for discovery and validation. Associations between asthma and neoplasms were discovered in data from the State of Florida. Validation was conducted on eight cohorts of patients with asthma, and seven subtypes of asthma and COPD using datasets from the State of California. Control groups were matched by gender, age, race, and history of tobacco use. Odds ratio analysis with Bonferroni-Holm correction measured the association of asthma and COPD with 26 different benign and malignant neoplasms. ICD9CM codes were used to identify exposures and outcomes.

**Findings:** CICT identified 17 associations between asthma and the risk of neoplasia in the discovery dataset. In the validation studies, 208 case-control analyses were conducted between subtypes of Asthma (N= 999,370, male= 33%, age= 50) and COPD (N=715,971, male = 50%, age=69) with the corresponding matched control groups (N=8,400,004, male= 42%, age= 47). Allergic asthma was associated with benign neoplasms of the meninges, salivary, pituitary, parathyroid, and thyroid glands (OR:1.52 to 2.52), and malignant neoplasms of the breast, intrahepatic biliary system, hematopoietic, and lymphatic system (OR: 1.45 to 2.05). COPD was associated with malignant neoplasms in the lung, bladder, and hematopoietic systems.

**Interpretation:** The combined use of machine learning methods for knowledge discovery and epidemiological methods shows that allergic asthma is associated with the development of neoplasia, including in glandular organs, ductal tissues, and hematopoietic systems. Also, our findings differentiate the pattern of neoplasms between allergic asthma and obstructive asthma. This suggests that inflammatory pathways that are active in asthma also contribute to neoplastic transformation in specific organ systems such as secretory organs.

**Funding:** **None**

**At a Glance Commentary:** Over the past three decades, studies have suggested that asthma could increase the risk of developing cancer, but a consensus has not been reached. The debate persists because the current evidence has been derived using cross-sectional statistical designs, limited datasets, and small cohorts and conflicting results. In addition, the mechanism by which allergic airway inflammation contributes to neoplastic transformation is postulated but not proven.

Here, we present the largest study to date on this association in patients with asthma or COPD. A knowledge discovery method was used for hypothesis generation that, when combined with epidemiological reasoning tools, identified associations between airway disease and neoplasia. The results reveal novel relationships between allergic asthma and benign glandular tumors and confirm the well-known connections between COPD and lung cancer. Further, we identified a novel association between COPD and asthma with hematological malignancies. These findings rectify contradictory results from other studies and demonstrate more specifically that the types of neoplasms associated with asthma compared to COPD that infers mechanistic plausibility.

## Introduction

Asthma is a common chronic disease that affects nearly 10% of the US population and is associated with an annual cost of approximately 80 billion dollars (1-3). Studies have demonstrated that the chronic systemic inflammation that is associated with asthma may contribute to the development and severity of comorbidities (4-6), including obesity, gastroesophageal reflux, psychological problems, chronic infections, and hormonal disturbances (7-10). Chronic inflammation is also known for its effect on neoplastic transformation and increased risk of cancer (11-13) through pathways that are also activated in asthma. Therefore, understanding the associations between asthma and neoplasia could inform both the pathobiology of chronic airway inflammation as well as the mechanisms that underlie the development of malignancy.

Definitive population-level evidence on the relationship between asthma and the development of neoplasia does not exist. A meta-analysis of 18 studies conducted between 1966 to 2002 suggested a 1.8 fold increase in lung cancer risk in patients with asthma (14) and a nationwide study in Sweden identified an increased incidence ratio of malignancies in 15 organ systems including leukemia, gastrointestinal, lung, prostate, nervous system, and thyroid gland with asthma (15). Another study on hospitalized patients in Sweden reported an increased incidence of all cancer, particularly stomach and colon cancer, in hospitalized asthma patients {Ji, 2009 #489}. In contrast, a longitudinal study of a US-based cohort (1982-2000) (17) found a protective effect of asthma on cancer mortality. Additional smaller-scale studies have also identified an association between asthma and increased risk of prostate cancer (18), hematopoietic malignancies (19), and a protective effect on adenocarcinoma of the pancreas. These studies are limited by small sample sizes, a lack of statistical power, biases in participant selection, exposure measurement (17, 20-24), and the limited type of neoplasms examined. Considering the significant clinical and economic impact that asthma and neoplasia have on society, additional studies need to be done to address these limitations and precisely determine if an association exists (17, 21).

To this end, we studied two large-scale longitudinal administrative datasets of over two hundred million hospitalization records in the US. We determined the association patterns between neoplasms and three subtypes of asthma and compared the results with four subtypes of Chronic Obstructive Pulmonary Disease (COPD). We developed a novel knowledge discovery method to identify disease-disease associations in high-dimensional data and validated the new findings with standard statistical methodology. We used a rigorous method for discovery and validation in one of the largest datasets of asthma and COPD analyzed to date. Our findings demonstrate that asthma is associated with a range of benign glandular neoplasms and malignant neoplasms in other organs, particularly secretory tissues.

## Materials and Methods

### Data Sources

The Healthcare Cost and Utilization Project (HCUP) is a federal-state-industry partnership sponsored by the Agency for Healthcare Research and Quality (25). Deidentified observational claim data from the HCUP was used for discovery and validation. The **S**tate **I**npatient **D**atabase (SID) and **E**mergency **D**epartment **D**atabase (EDD) from the state of Florida from 2004 to 2014 were used for exploratory analyses (25). Validation was conducted using data from the California HCUP SID and EDD from 2005 to 2011(25). HCUP data is a census of all discharges (26), and SID contains the records for all-payers, including the uninsured, comprising all non-federal acute care hospitals in participating states (27), and together, about 95 percent of all US community hospital discharges (28, 29) (30). The ED contains all visits to the affiliated emergency department that did not result in a hospitalization. Each HCUP record represents a patient encounter and includes demographic data, clinical diagnoses, comorbidities, procedures, total costs of hospitalization and other information from claim records. HCUP also has an identifier that can connect all encounters across the SID and ED to create a longitudinal record for each patient. The identifier, ‘VisitLink,’ is assigned through a verification process that ensures the correct assignment to all admissions of a patient (31).

The study population included patients 18 years or older who had at least two inpatient or emergency department observations in our dataset. The hypothesis generation method employed in the discovery phase, CICT, uses the diagnoses that emerge between visits to predict potential causal associations. Accordingly, we included only patients with more than one visit. Patients were excluded if the VisitLink information, a revalidated patient identifier, was missing or if the patient died during hospitalization. The CONSORT diagram in Figure 1 summarizes the data preparation process, and population characteristics are shown in Table 1 (31).

**Table 1:** Basic characteristics of patients with asthma and COPD by subtype - California HCUP SID, EDD 2005-2011.

| Characteristics, N (%) | Population Data | Asthma Subtypes |  |  | Asthma (493.*) | COPD Subtypes |  |  |  | COPD (490, 491, 492, 494, 495, 496) |
| --- | --- | --- | --- | --- | --- | --- | --- | --- | --- | --- |
|  |  | Extrinsic asthma (493.0) | Chronic obstructive asthma (493.2) | Asthma, unspecified (493.9) |  | Chronic bronchitis (491) | Emphysema (492) | Bronchiectasis (494) | Chronic airway obstruction, not elsewhere classified. (496) |  |
| **Total count** | 12,188,304 | 28149 (0.33%) | 235446 (2.73%) | 853556 (9.22%) | 999370 (10.63%) | 390862 (4.45%) | 97956 (1.15%) | 29539 (0.35%) | 715957 (7.85%) | 715971 (6.95%) |
| **Age (SD)** | 42.65 (23.86) | 47 (18) | 65 (15) | 48 (19) | 50 (20) | 68 (15) | 68 (14) | 71 (15) | 69 (14) | 69 (14) |
| **Sex** |  |  |  |  |  |  |  |  |  |  |
| **Men** | 5,314,100 (43.6%) | 8864 (31.49%) | 88982 (37.79%) | 269308 (31.55%) | 329413 (32.96%) | 186542 (47.73%) | 52571 (53.67%) | 11610 (39.30%) | 361018 (50.42%) | 361027 (50.42%) |
| **Women** | 6,795,494 (56.4%) | 19285 (68.51%) | 146464 (62.21%) | 584248 (68.45%) | 669957 (67.04%) | 204320 (52.27%) | 45385 (46.33%) | 17929 (60.70%) | 354939 (49.58%) | 354944 (49.58%) |
| **Race** |  |  |  |  |  |  |  |  |  |  |
| **White** | 6,194,004 (51.4%) | 16392 (58.23%) | 152944 (64.96%) | 474326 (55.57%) | 574255 (57.46%) | 274497 (70.23%) | 74496 (76.05%) | 18864 (63.86%) | 520076 (72.64%) | 519785 (72.60%) |
| **Black** | 1,160,655 (9.6%) | 3766 (13.38%) | 28163 (11.96%) | 120800 (14.15%) | 134259 (13.43%) | 37209 (9.52%) | 8345 (8.52%) | 1495 (5.06%) | 60958 (8.51%) | 60616 (8.47%) |
| **Hispanic** | 3,468,709 (28.8%) | 5194 (18.45%) | 32482 (13.80%) | 183721 (21.52%) | 202252 (20.24%) | 48715 (12.46%) | 9003 (9.19%) | 4217 (14.28%) | 84978 (11.87%) | 85020 (11.87%) |
| **Asian or Pacific Islander** | 807,814 (6.7%) | 2003 (7.12%) | 16926 (7.19%) | 51158 (5.99%) | 61992 (6.20%) | 22847 (5.85%) | 4468 (4.56%) | 4320 (14.62%) | 36803 (5.14%) | 37095 (5.18%) |
| **Native American** | 27,493 (0.2%) | 75 (0.27%) | 585 (0.25%) | 2467 (0.29%) | 2771 (0.28%) | 764 (0.20%) | 167 (0.17%) | 36 (0.12%) | 1322 (0.18%) | 1535 (0.21%) |
| **Others** | 389,883 (3.2%) | 719 (2.55%) | 4346 (1.85%) | 21084 (2.47%) | 23841 (2.39%) | 6830 (1.75%) | 1477 (1.51%) | 607 (2.05%) | 11820 (1.65%) | 11920 (1.66%) |
| **Smoking** | 1,664,159 (13.8%) | 5762 (20.47%) | 88443 (37.56%) | 212813 (24.93%) | 262847 (26.30%) | 173756 (44.45%) | 44051 (44.97%) | 4361 (14.76%) | 283411 (39.58%) | 283423 (39.59%) |

**Figure 1:**
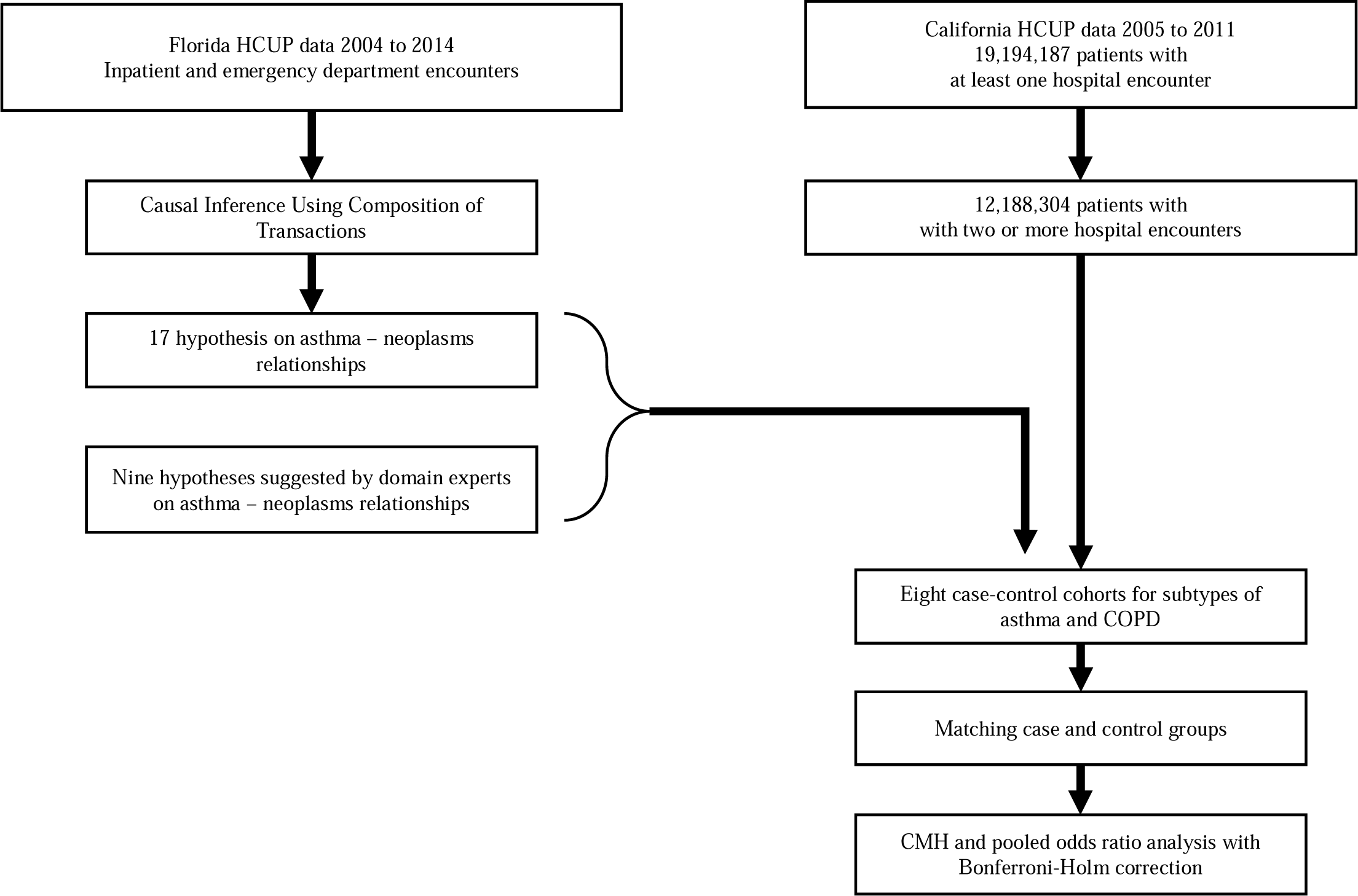
CONSORT diagram.

### Discovery analysis

Evaluation of all associations between subtypes of asthma and neoplasia could be computationally prohibitive and result in a significant Type I error. Accordingly, to limit the number of comparisons, we developed the Causal Inference using the Composition of Transitions (CICT) method (32). CICT assigns the primary diagnosis of each encounter as the main clinical condition of the patient and assumes that a patient’ s two separate encounters for different primary diagnoses is a transition from the first clinical condition to the second. Here, CICT uses longitudinal data from the population to count all such transitions among patients during a specific time period. The information that results from this algorithm is a Poisson stochastic process. CICT reasons that in such a Poisson process, the frequency of transition between each pair of random events should be proportional to the frequency of the first and the second condition. A non-random event (e.g., causal) would appear as a disturbance in the default Poisson process of events because there is an increased rate of transition to specific conditions (effects) compared to a reduced rate of transition to random events. For example, since COPD is associated with a risk of lung cancer, there are higher rates of transition to lung cancer, and proportionally lower rates to other unrelated conditions. CICT uses supervised machine learning methods and a set of labeled causal and random transitions to learn patterns that discriminate random versus causal transitions (33). Then, CICT uses the learned patterns to predict relationships among various clinical conditions. For this study, we applied CICT to a discovery dataset from the state of Florida to identify relationships between asthma and 514 different types of benign and malignant neoplasia. The clinical conditions were identified using the ICD9CM code of primary diagnosis for admission. Table 2 shows the predicted relationships between asthma and neoplasia, and Figure 2 is the CICT output graph network of the predicted connections of asthma and neoplasms of different organs.

**Table 2:** CICT generated predicted strength for the hypothesis on the effect of subtypes of asthma (columns) on inducing neoplasms (rows) using Florida 2004-2014 SID and ED data. The number represents the predicted strength of hypotheses, range 0-1; 1 indicating a strong probability, 0, indicating a random relationship. The 95% confidence interval of each prediction is given in the parenthesis. Malignant neoplasms are abbreviated to MN and benign neoplasms to BN.

| ICD9CM code | Exposure |  | Chronic obstructive asthma, unspecified (ICD:493.2) |
| --- | --- | --- | --- |
|  | Event | Asthma, unspecified type, with (acute) exacerbation (ICD: 493.9) |  |
| 210.2 | BN of major salivary glands | 0.93 (0.81 - 1.00) | 0.92 (0.81 - 1.00) |
| 227.3 | BN of pituitary gland and craniopharyngeal duct | 0.73 (0.57 - 0.89) | 0.73 (0.63 - 0.83) |
| 227.1 | BN of parathyroid gland | 0.83 (0.72 - 0.94) | 0.86 (0.75 - 0.97) |
| 225.2 | BN of cerebral meninges | 0.65 (0.28 - 1.00) | 0.57 (0.33 - 0.81) |
| 226 | BN of thyroid glands | 0.87 (0.76 - 0.98) | 0.90 (0.79 - 1.00) |
| 153.4 | MN of cecum | N/A | 0.63 (0.43 - 0.83) |
| 153.6 | MN of ascending colon | 0.66 (0.55 - 0.77) | 0.56 (0.44 - 0.67) |
| 153.9 | MN of colon, unspecified site | N/A | 0.80 (0.70 - 0.90) |
| 153.0 | MN of hepatic flexure | N/A | 0.94 (0.83 - 1.00) |
| 153.1 | MN of transverse colon | 0.89 (0.78 - 1.00) | 0.82 (0.72 - 0.93) |
| 155.1 | MN of intrahepatic bile ducts | N/A | 0.94 (0.83 - 1.00) |
| 156.1 | MN of extrahepatic bile ducts | N/A | 0.93 (0.82 - 1.00) |
| 174.4 | MN of upper-outer quadrant of female breast | 0.74 (0.64 - 0.84) | 0.73 (0.63 - 0.83) |
| 162.4 | MN of middle lobe, bronchus or lung | N/A | 0.96 (0.85 - 1.00) |
| 188.9 | MN of bladder, part unspecified | 0.78<br>(0.56 - 1.00) | 0.69 (0.59 - 0.80) |
| 188.2 | MN of lateral wall of the urinary bladder | N/A | 0.96 (0.85 - 1.00) |
| 238.7 | Other lymphatic and hematopoietic tissues | 0.74 (0.63 - 0.86) | 0.69 (0.59 - 0.80) |

**Figure 2:**
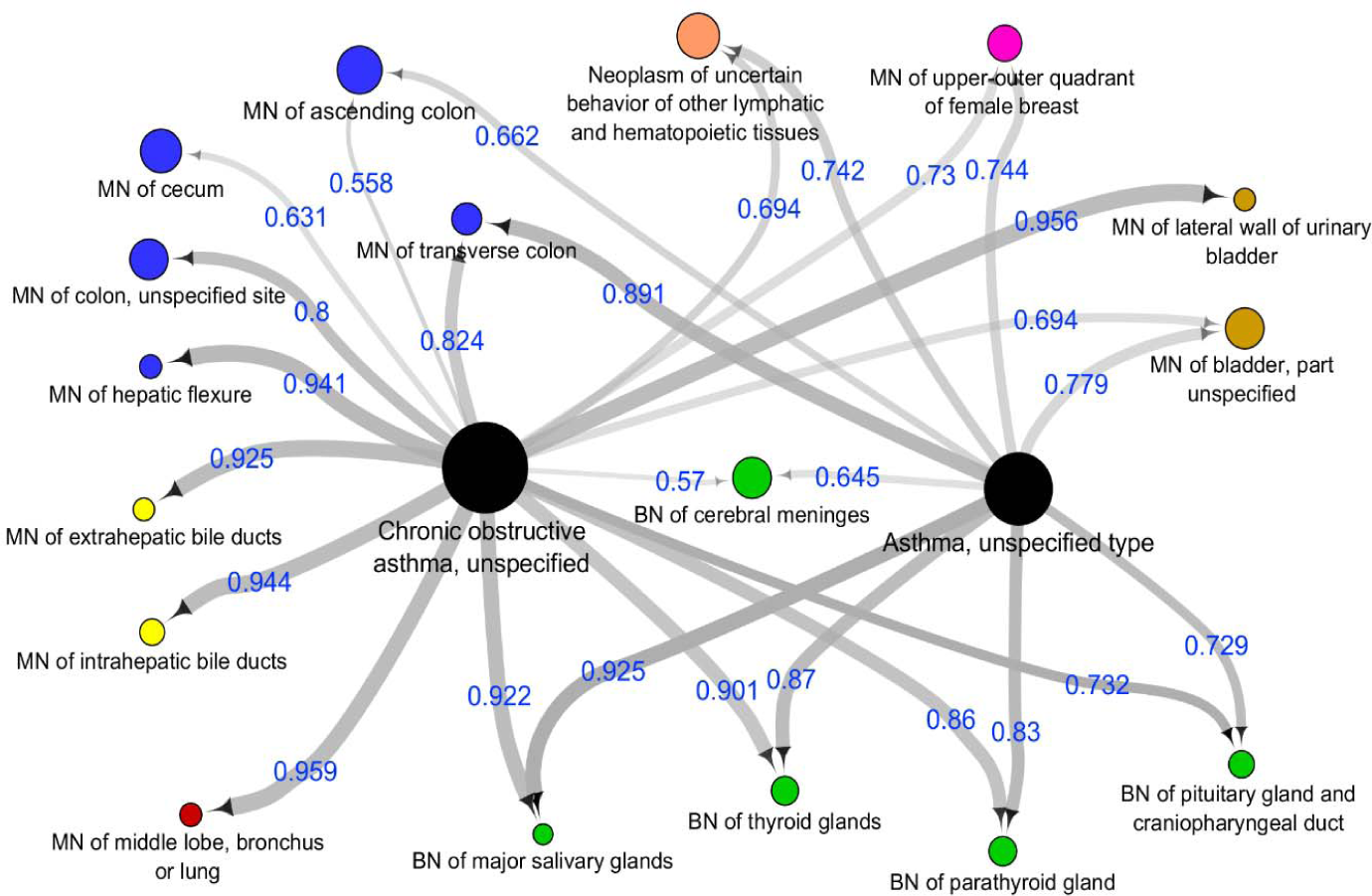
CICT generated relationships between asthma and neoplasia using Florida HCUP 2004-2014 data. Edge values are CICT predicted strength for the hypothesis (0 to 1). Nodes are color-coded and organized based on their clinical relevance: asthma subtypes (center, black), MN of gastrointestinal (left-top, blue), MN of bile ducts(left, yellow), MN or lung (left-bottom, red), BN of cerebral meninges and glands (bottom, green), MN urinary track (right, brown), MN of lymphatic and hematopoietic-top (top, orange), MN of the upper outer breast (top, pink). Node size is non-linearly proportional to the patients’ frequency. MN= Malignant neoplasm, BN= benign neoplasm.

### Validation analysis

To validate the hypotheses that were discovered in the CICT analysis, we conducted Odds Ratio (OR) analysis on the HCUP dataset from California. Exposures and outcome events were identified using ICD9CM codes for the primary diagnosis and up to 24 comorbidities. Race was stratified into six categories: Black, White, Hispanic, Asian or Pacific Islander, Native American, and others in the HCUP data. History of smoking was identified using the ICD9CM codes including, 305.1 (tobacco use disorder), 649.0 (tobacco use disorder complicating pregnancy, childbirth, or the puerperium), v15.82 (personal history of tobacco use), and 989.84 (toxic effect of tobacco). To discriminate the effect of asthma compared to COPD on the associations with neoplasms, we also performed OR analysis on subtypes of COPD with the same neoplasms. Descriptive statistics of individual characteristics at baseline were calculated for the whole population, asthma, and COPD subtype cohorts (Table 1). We computed frequencies for categorical variables and means with standard deviations for continuous variables. The significance threshold was established at P ≤ 0.05. All statistical analyses were performed using R 3.5.3 (R Foundation, Switzerland).

Each cohort of asthma and COPD was matched to an unaffected control cohort by race, gender, age, and smoking status using coarsened exact matching (CEM) (34, 35). CEM first transforms levels of continuous or categorical variables into a smaller number of categories and then conducts exact matching on the coarsened categories, while containing balancing error under the desired threshold. CEM makes no assumptions about the data generation process (beyond the usual ignorability assumptions)(35) and has been shown to yield estimates of causal effects that are lowest in variance and bias for any given sample size (36) which thus is preferred over matching by modeling (e.g., propensity score). The remaining imbalance and heterogeneity between case and control groups were assessed using the multivariate standardized mean difference (SMD) metric.

The association between asthma and neoplasia was estimated using both the Cochran-Mantel-Haenszel (CMH) (37, 38) common odds ratio (cOR) and pooled odds ratio over the whole cohort. cOR is a weighted aggregation of OR.s in balanced strata of the population and is considered a more accurate estimate of association than pooled OR, which is a raw, and often higher, estimate of association in the cohort (Appendix 4). We report cOR and 95% confidence intervals (CIs) for 26 neoplasms (column 1 of Table 3) in the eight matched case-control subtypes of asthma and COPD (row 1 of Table 3). A Bonferroni-Holm (39) correction was applied to each asthma and COPD subtype cohort to control the family-wise error rate.

**Table 3:**
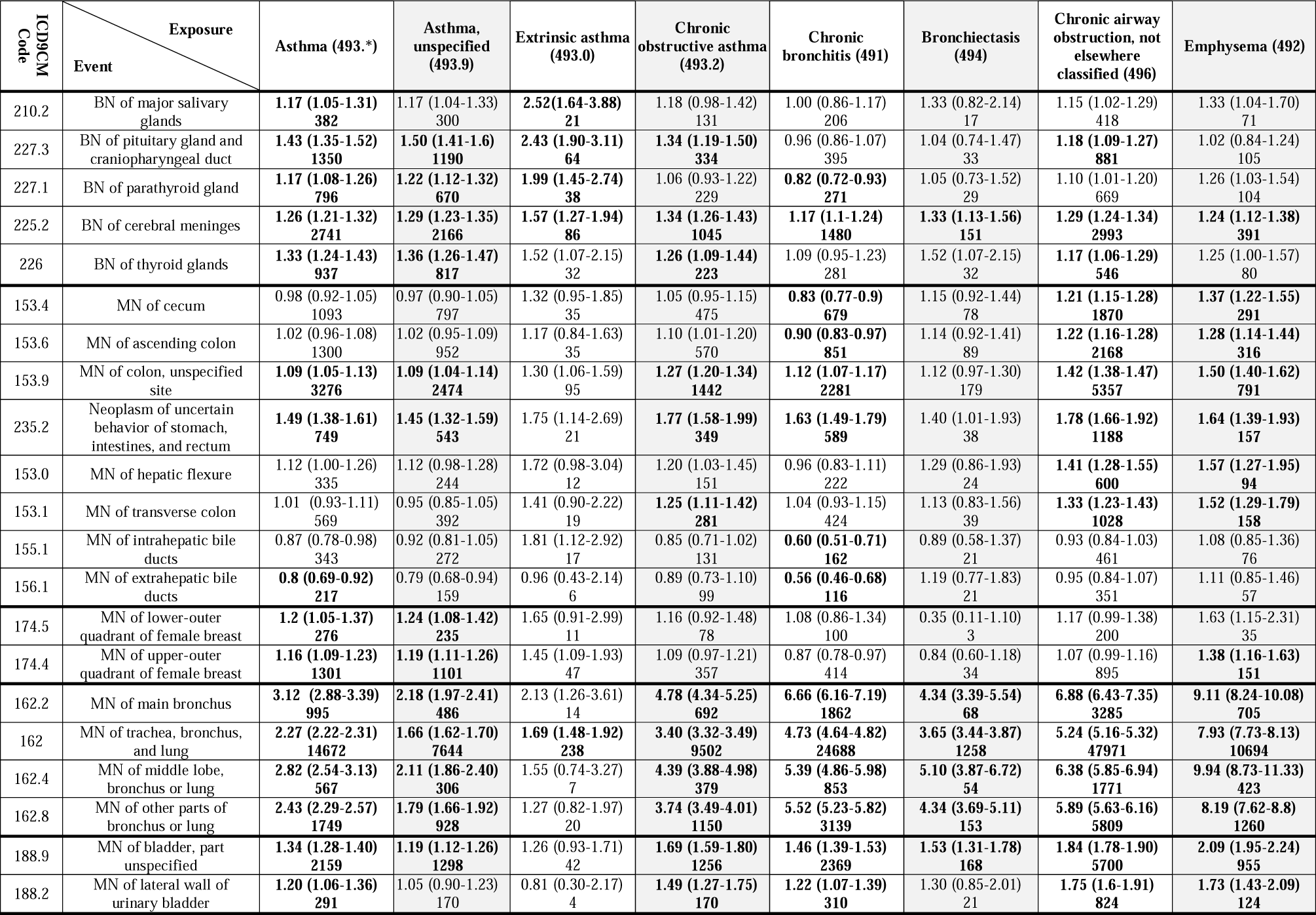
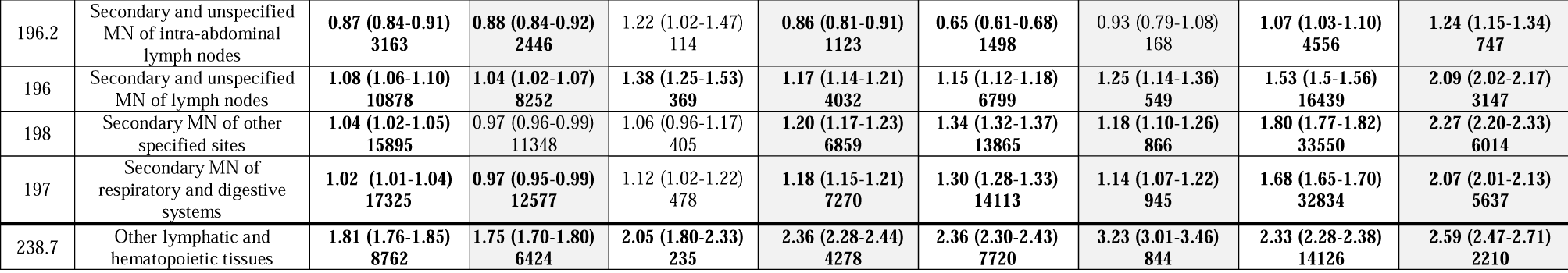
Cochran-Mantel-Haenszel (CMH) common odds ratios of neoplasms in subtypes of asthma and COPD compared to controls. Values in parenthesis show the lower and upper bounds of the 95% confidence interval. In each cell, the second line shows the prevalence of each outcome in each exposure. Bold type indicates statistically significant odds ratios when the Bonferroni-Holm adjusted p-values are less than 0.05. In event description, BN and MN correspondingly stand for ‘benign neoplasm’ and ‘malignant neoplasm.’

| ICD9CM Code | Exposure<br>Event | Asthma (493.*) | Asthma, unspecified (493.9) | Extrinsic asthma (493.0) | Chronic obstructive asthma (493.2) | Chronic bronchitis (491) | Bronchiectasis (494) | Chronic airway obstruction, not elsewhere classified (496) | Emphysema (492) |
| --- | --- | --- | --- | --- | --- | --- | --- | --- | --- |
| 210.2 | BN of major salivary glands | **1.17 (1.05-1.31)**<br>382 | 1.17 (1.04-1.33)<br>300 | **2.52 (1.64-3.88)**<br>21 | 1.18 (0.98-1.42)<br>131 | 1.00 (0.86-1.17)<br>206 | 1.33 (0.82-2.14)<br>17 | 1.15 (1.02-1.29)<br>418 | 1.33 (1.04-1.70)<br>71 |
| 227.3 | BN of pituitary gland and craniopharyngeal duct | **1.43 (1.35-1.52)**<br>1350 | **1.50 (1.41-1.6)**<br>1190 | **2.43 (1.90-3.11)**<br>64 | **1.34 (1.19-1.50)**<br>334 | 0.96 (0.86-1.07)<br>395 | 1.04 (0.74-1.47)<br>33 | **1.18 (1.09-1.27)**<br>881 | 1.02 (0.84-1.24)<br>105 |
| 227.1 | BN of parathyroid gland | **1.17 (1.08-1.26)**<br>796 | **1.22 (1.12-1.32)**<br>670 | **1.99 (1.45-2.74)**<br>38 | 1.06 (0.93-1.22)<br>229 | **0.82 (0.72-0.93)**<br>271 | 1.05 (0.73-1.52)<br>29 | 1.10 (1.01-1.20)<br>669 | 1.26 (1.03-1.54)<br>104 |
| 225.2 | BN of cerebral meninges | **1.26 (1.21-1.32)**<br>2741 | **1.29 (1.23-1.35)**<br>2166 | **1.57 (1.27-1.94)**<br>86 | **1.34 (1.26-1.43)**<br>1045 | **1.17 (1.1-1.24)**<br>1480 | **1.33 (1.13-1.56)**<br>151 | **1.29 (1.24-1.34)**<br>2993 | **1.24 (1.12-1.38)**<br>391 |
| 226 | BN of thyroid glands | **1.33 (1.24-1.43)**<br>937 | **1.36 (1.26-1.47)**<br>817 | 1.52 (1.07-2.15)<br>32 | **1.26 (1.09-1.44)**<br>223 | 1.09 (0.95-1.23)<br>281 | 1.52 (1.07-2.15)<br>32 | **1.17 (1.06-1.29)**<br>546 | 1.25 (1.00-1.57)<br>80 |
| 153.4 | MN of cecum | 0.98 (0.92-1.05)<br>1093 | 0.97 (0.90-1.05)<br>797 | 1.32 (0.95-1.85)<br>35 | 1.05 (0.95-1.15)<br>475 | **0.83 (0.77-0.9)**<br>679 | 1.15 (0.92-1.44)<br>78 | **1.21 (1.15-1.28)**<br>1870 | **1.37 (1.22-1.55)**<br>291 |
| 153.6 | MN of ascending colon | 1.02 (0.96-1.08)<br>1300 | 1.02 (0.95-1.09)<br>952 | 1.17 (0.84-1.63)<br>35 | 1.10 (1.01-1.20)<br>570 | **0.90 (0.83-0.97)**<br>851 | 1.14 (0.92-1.41)<br>89 | **1.22 (1.16-1.28)**<br>2168 | **1.28 (1.14-1.44)**<br>316 |
| 153.9 | MN of colon, unspecified site | **1.09 (1.05-1.13)**<br>3276 | **1.09 (1.04-1.14)**<br>2474 | 1.30 (1.06-1.59)<br>95 | **1.27 (1.20-1.34)**<br>1442 | **1.12 (1.07-1.17)**<br>2281 | 1.12 (0.97-1.30)<br>179 | **1.42 (1.38-1.47)**<br>5357 | **1.50 (1.40-1.62)**<br>791 |
| 235.2 | Neoplasm of uncertain behavior of stomach, intestines, and rectum | **1.49 (1.38-1.61)**<br>749 | **1.45 (1.32-1.59)**<br>543 | 1.75 (1.14-2.69)<br>21 | **1.77 (1.58-1.99)**<br>349 | **1.63 (1.49-1.79)**<br>589 | 1.40 (1.01-1.93)<br>38 | **1.78 (1.66-1.92)**<br>1188 | **1.64 (1.39-1.93)**<br>157 |
| 153.0 | MN of hepatic flexure | 1.12 (1.00-1.26)<br>335 | 1.12 (0.98-1.28)<br>244 | 1.72 (0.98-3.04)<br>12 | 1.20 (1.03-1.45)<br>151 | 0.96 (0.83-1.11)<br>222 | 1.29 (0.86-1.93)<br>24 | **1.41 (1.28-1.55)**<br>600 | **1.57 (1.27-1.95)**<br>94 |
| 153.1 | MN of transverse colon | 1.01 (0.93-1.11)<br>569 | 0.95 (0.85-1.05)<br>392 | 1.41 (0.90-2.22)<br>19 | **1.25 (1.11-1.42)**<br>281 | 1.04 (0.93-1.15)<br>424 | 1.13 (0.83-1.56)<br>39 | **1.33 (1.23-1.43)**<br>1028 | **1.52 (1.29-1.79)**<br>158 |
| 155.1 | MN of intrahepatic bile ducts | 0.87 (0.78-0.98)<br>343 | 0.92 (0.81-1.05)<br>272 | 1.81 (1.12-2.92)<br>17 | 0.85 (0.71-1.02)<br>131 | **0.60 (0.51-0.71)**<br>162 | 0.89 (0.58-1.37)<br>21 | 0.93 (0.84-1.03)<br>461 | 1.08 (0.85-1.36)<br>76 |
| 156.1 | MN of extrahepatic bile ducts | **0.8 (0.69-0.92)**<br>217 | 0.79 (0.68-0.94)<br>159 | 0.96 (0.43-2.14)<br>6 | 0.89 (0.73-1.10)<br>99 | **0.56 (0.46-0.68)**<br>116 | 1.19 (0.77-1.83)<br>21 | 0.95 (0.84-1.07)<br>351 | 1.11 (0.85-1.46)<br>57 |
| 174.5 | MN of lower-outer quadrant of female breast | **1.2 (1.05-1.37)**<br>276 | **1.24 (1.08-1.42)**<br>235 | 1.65 (0.91-2.99)<br>11 | 1.16 (0.92-1.48)<br>78 | 1.08 (0.86-1.34)<br>100 | 0.35 (0.11-1.10)<br>3 | 1.17 (0.99-1.38)<br>200 | 1.63 (1.15-2.31)<br>35 |
| 174.4 | MN of upper-outer quadrant of female breast | **1.16 (1.09-1.23)**<br>1301 | **1.19 (1.11-1.26)**<br>1101 | 1.45 (1.09-1.93)<br>47 | 1.09 (0.97-1.21)<br>357 | 0.87 (0.78-0.97)<br>414 | 0.84 (0.60-1.18)<br>34 | 1.07 (0.99-1.16)<br>895 | **1.38 (1.16-1.63)**<br>151 |
| 162.2 | MN of main bronchus | **3.12 (2.88-3.39)**<br>995 | **2.18 (1.97-2.41)**<br>486 | 2.13 (1.26-3.61)<br>14 | **4.78 (4.34-5.25)**<br>692 | **6.66 (6.16-7.19)**<br>1862 | **4.34 (3.39-5.54)**<br>68 | **6.88 (6.43-7.35)**<br>3285 | **9.11 (8.24-10.08)**<br>705 |
| 162 | MN of trachea, bronchus, and lung | **2.27 (2.22-2.31)**<br>14672 | **1.66 (1.62-1.70)**<br>7644 | **1.69 (1.48-1.92)**<br>238 | **3.40 (3.32-3.49)**<br>9502 | **4.73 (4.64-4.82)**<br>24688 | **3.65 (3.44-3.87)**<br>1258 | **5.24 (5.16-5.32)**<br>47971 | **7.93 (7.73-8.13)**<br>10694 |
| 162.4 | MN of middle lobe, bronchus or lung | **2.82 (2.54-3.13)**<br>567 | **2.11 (1.86-2.40)**<br>306 | 1.55 (0.74-3.27)<br>7 | **4.39 (3.88-4.98)**<br>379 | **5.39 (4.86-5.98)**<br>853 | **5.10 (3.87-6.72)**<br>54 | **6.38 (5.85-6.94)**<br>1771 | **9.94 (8.73-11.33)**<br>423 |
| 162.8 | MN of other parts of bronchus or lung | **2.43 (2.29-2.57)**<br>1749 | **1.79 (1.66-1.92)**<br>928 | 1.27 (0.82-1.97)<br>20 | **3.74 (3.49-4.01)**<br>1150 | **5.52 (5.23-5.82)**<br>3139 | **4.34 (3.69-5.11)**<br>153 | **5.89 (5.63-6.16)**<br>5809 | **8.19 (7.62-8.8)**<br>1260 |
| 188.9 | MN of bladder, part unspecified | **1.34 (1.28-1.40)**<br>2159 | **1.19 (1.12-1.26)**<br>1298 | 1.26 (0.93-1.71)<br>42 | **1.69 (1.59-1.80)**<br>1256 | **1.46 (1.39-1.53)**<br>2369 | **1.53 (1.31-1.78)**<br>168 | **1.84 (1.78-1.90)**<br>5700 | **2.09 (1.95-2.24)**<br>955 |
| 188.2 | MN of lateral wall of urinary bladder | **1.20 (1.06-1.36)**<br>291 | 1.05 (0.90-1.23)<br>170 | 0.81 (0.30-2.17)<br>4 | **1.49 (1.27-1.75)**<br>170 | **1.22 (1.07-1.39)**<br>310 | 1.30 (0.85-2.01)<br>21 | **1.75 (1.6-1.91)**<br>824 | **1.73 (1.43-2.09)**<br>124 |
| 196.2 | Secondary and unspecified<br>MN of intra-abdominal<br>lymph nodes | **0.87 (0.84-0.91)**<br>**3163** | **0.88 (0.84-0.92)**<br>**2446** | 1.22 (1.02-1.47)<br>114 | **0.86 (0.81-0.91)**<br>**1123** | **0.65 (0.61-0.68)**<br>**1498** | 0.93 (0.79-1.08)<br>168 | **1.07 (1.03-1.10)**<br>**4556** | **1.24 (1.15-1.34)**<br>**747** |
| 196 | Secondary and unspecified<br>MN of lymph nodes | **1.08 (1.06-1.10)**<br>**10878** | **1.04 (1.02-1.07)**<br>**8252** | **1.38 (1.25-1.53)**<br>**369** | **1.17 (1.14-1.21)**<br>**4032** | **1.15 (1.12-1.18)**<br>**6799** | **1.25 (1.14-1.36)**<br>**549** | **1.53 (1.5-1.56)**<br>**16439** | **2.09 (2.02-2.17)**<br>**3147** |
| 198 | Secondary MN of other<br>specified sites | **1.04 (1.02-1.05)**<br>**15895** | 0.97 (0.96-0.99)<br>11348 | 1.06 (0.96-1.17)<br>405 | **1.20 (1.17-1.23)**<br>**6859** | **1.34 (1.32-1.37)**<br>**13865** | **1.18 (1.10-1.26)**<br>**866** | **1.80 (1.77-1.82)**<br>**33550** | **2.27 (2.20-2.33)**<br>**6014** |
| 197 | Secondary MN of<br>respiratory and digestive<br>systems | **1.02 (1.01-1.04)**<br>**17325** | **0.97 (0.95-0.99)**<br>**12577** | 1.12 (1.02-1.22)<br>478 | **1.18 (1.15-1.21)**<br>**7270** | **1.30 (1.28-1.33)**<br>**14113** | **1.14 (1.07-1.22)**<br>**945** | **1.68 (1.65-1.70)**<br>**32834** | **2.07 (2.01-2.13)**<br>**5637** |
| 238.7 | Other lymphatic and<br>hematopoietic tissues | **1.81 (1.76-1.85)**<br>**8762** | **1.75 (1.70-1.80)**<br>**6424** | **2.05 (1.80-2.33)**<br>**235** | **2.36 (2.28-2.44)**<br>**4278** | **2.36 (2.30-2.43)**<br>**7720** | **3.23 (3.01-3.46)**<br>**844** | **2.33 (2.28-2.38)**<br>**14126** | **2.59 (2.47-2.71)**<br>**2210** |

## Results

Four cohorts were evaluated for asthma and its three subtypes based on ICD-9-CM codes: 493.*: combined asthma (N= 999370), extrinsic asthma (N= 28149, male= 31%, age= 47), chronic obstructive asthma (N= 235446, male= 38%, age= 65), and asthma unspecified (N= 853556, male= 32%, age= 48). We excluded intrinsic asthma groups due to the low number of exposed patients (N= 1167, 0.12% of the asthma patients) and because few or no neoplasms of interest were identified in both case and control groups. Four cohorts were designed for COPD and relevant conditions: chronic bronchitis (N= 390862, male= 48%, age=68), emphysema (N= 97956, male= 54%, age= 68), bronchiectasis (N= 29539, male= 39%, age= 71), and chronic airway obstruction, not elsewhere classified (N= 715957, male= 50%, age= 69). COPD patients were older and equally distributed between both sex groups and asthma patients were predominantly middle-aged females. The highest rate of smoking was observed in patients with emphysema (45%) and the lowest in allergic asthma (20%).

### CICT identified associations between asthma and neoplasms

Among 524 neoplasms categorized by ICD9CM codes, CICT identified 12 malignant and five benign neoplasms that were associated with asthma (Table 2, Figure 2). The strongest relationship was predicted between chronic obstructive asthma (ICD9:493.2) and malignant neoplasm of the middle lobe, bronchus, or lung (ICD9CM:162.4, estimate: 0.96, 95% CI: 0.85 - 1.00). Other associations with asthma were malignant neoplasm of transverse colon (ICD9CM 153.1, 0.82, CI:0.72 - 0.93), bile ducts(ICD9CM 155.1, 0.94, CI:0.83 - 1.00), bladder (ICD9CM 188.9, 0.78, CI: 0.56 - 1.00) and lymphatic and hematopoietic tissues (ICD9CM 238.7, 0.74, CI:0.63 - 0.86). The strongest relationship between asthma and benign neoplasms was with benign neoplasms of salivary glands (ICD9CM 210.2, 0.93, CI: 0.81 - 1.00), followed by benign glandular neoplasms of the parathyroid (ICD9CM 227.1, 0.86, CI: 0.75 - 0.97), thyroid (ICD9CM 226, 0.90, CI: 0.79 - 1.00) and pituitary glands (ICD9CM 227.3, 0.73, CI:0.63 - 0.83). Therefore, CICT analysis demonstrated a relationship between asthma and the development of neoplasia of glandular structures, the gastrointestinal system, biliary ducts, lymphatic and hematopoietic systems, and breast.

### Validation of CICT identified associations

The association between asthma and 26 neoplasia were evaluated in the four case-control cohorts as described in *Validation analysis*. We also evaluated the associations between neoplasia and COPD and related conditions in four additional case-control cohorts, to contrast differences in associations between asthma compared to COPD. In all cohorts, the application of matching reduced standardized mean difference (SMD) of the case and the matched control groups to 0.0 indicating the case and matched cohorts in 208 different studies have similar demographic features and smoking exposure. (34)

Approximately 8.4 million matched controls were included in each cohort: asthma(N= 8,370,574), extrinsic asthma (N= 8,398,723), chronic obstructive asthma (N = 8,400,004), unspecified asthma (N = 8,400,004), chronic bronchitis (N = 8,400,004), emphysema (N = 8,397,079), bronchiectasis (N = 8,399,392) chronic airway obstruction (N = 8,400,004). There was a history of asthma in 10.63 %,and COPD in 6.95% of the total included population, consistent with national averages (3) (1) (40). On average, patients with chronic obstructive asthma and COPD tended to be older than patients with other asthma subtypes. Patients with extrinsic asthma, asthma unspecified, and bronchiectasis were less likely to be former smokers than patients with other subtypes of asthma and COPD. Bronchiectasis and subtypes of asthma were more likely to be women than patients with chronic bronchitis, emphysema, and chronic airway obstruction - not elsewhere classified. Compared with healthy people with no history of asthma or COPD, patients with asthma were more likely to be older adults, females, white or black race, and smokers. A total of 139,038 incident cases of neoplasms were identified among patients with asthma, 338,418 in those exposed to COPD, and 518,138 in the control cohort. Age, sex, race, and smoking distribution in the validation cohorts are shown in Table 1.

The cOR for asthma, COPD, and their subtypes compared to the control cohort for individual cancers are presented in Table 3 and Appendix Figures 2 and Figure 3. The balancing, conducted by CEM, ensures the similarity of potential confounders between cases and controls in each stratum. To provide a basis for comparison, pooled ORs are presented in Appendix Table 1. Extrinsic asthma was associated with benign neoplasms of the salivary, parathyroid, thyroid, and pituitary and craniopharyngeal duct (cOR:1.52-2.52); and with malignant neoplasms of intrahepatic bile ducts and upper-outer quadrant of female breast (cOR 1.45-1.81), all confirming the results of CICT knowledge discovery method. Among subtypes of asthma, obstructive asthma had the highest risk of various pulmonary cancers (cOR: 3.40 to 4.78), compared with extrinsic asthma, which showed a weaker and often insignificant association with lung cancers. Emphysema and Chronic airway obstruction, not elsewhere classified (496) had the strongest associations with primary malignant neoplasms (cOR 1.07 – 9.94). Also, emphysema showed the strongest association with lung cancers (cOR 7.93-9.94). (table 3 and Appendix figure 3). Therefore the associations between asthma and neoplasia that were identified using CICT and validated were benign neoplasms of the salivary, pituitary, parathyroid, and thyroid glands, and cerebral meninges and malignant neoplasms of the colon, stomach, breast, bronchus, and lung.

Subtypes of Asthma and COPD showed a weak association with secondary or metastatic neoplasms (Table 3 and Appendix figure 2). Emphysema was associated with increased odds for secondary malignant neoplasm of other specified sites (ICD9CM: 198, OR: 2.27, CI: 2.20-2.33), secondary malignant neoplasm of respiratory and digestive systems (ICD9CM: 197, OR: 2.07, CI: 2.01-2.13) and secondary malignancy of intra-abdominal lymph nodes (ICD9CM: 196.2, OR: 1.24, CI: 1.15-1.34). Subtypes of COPD are associated with hematopoietic and lymphatic cancers (ICD9CM: 238.7, OR: 2.33-3.23). Extrinsic asthma is also associated with increased odds of lymphatic and hematopoietic cancers (OR: 2.05, CI: 1.80-2.33).

## Discussion

The long-standing debate about the relationship between asthma and the development of neoplasia has been hampered by methodologic limitations and the inability to generate population-level evidence to properly examine the association. To overcome this barrier, we analyzed one of the largest datasets available of patients with asthma and COPD, approximating a population-level study of the association. Using a novel knowledge discovery method, CICT, and conventional statistical methods for validation, we confirmed associations between asthma, COPD and malignancy and identified associations between asthma and benign neoplasia that are consistent with the hypothesis that the chronic inflammation associated with asthma contributes to neoplastic transformation in the lung and other organs. This approach and the large size of the cohorts allowed us to assess associations with more strength than previous studies. Moreover, this is the first study that suggests that asthma could increase the risk of both benign and malignant neoplasia.

An important aspect of this study is the robust two-step design and the use of predictive, instead of inferential, modeling to find potential associations. In the first step, we deployed CICT, a machine learning knowledge discovery method to directly identify potential associations. CICT uses anomalies in a default Poisson process of patients’ transitions between clinical conditions to predict potential causal associations. Defining causal associations as an anomaly in the Poisson process of transitions is an innovative approach. Also, the use of predictive modeling to identify causal associations is a departure from standard methods that infer causality using mathematical or statistical inferential models to estimate the effect size after removing confounding effects. Notably, CICT does not use any contextual variable, such as age, gender, or comorbid conditions. In the second step, we used two independent large-scale claims dataset and standard statistical methods, controlling for confounders, to validate the hypotheses that were generated by CICT analysis in the first step. Combined, the use of knowledge discovery methods and validation by standard methods helped to draw a concise picture of the associations between asthma and the development of neoplasia. This shows that adding causality prediction to the analysis of big data can be used to identify relationships and interactions between diseases in a manner as close to population-level as possible.

The contrast between allergic and chronic obstructive asthma was critical to understanding the identification of associations with neoplasms. Chronic obstructive asthma, a code that includes patients with allergic and obstructive manifestations of asthma, showed similarities to both asthma and COPD but was associated with a pattern of neoplasia that overlapped more with COPD than asthma. This may be related to limitations in disease specificity and could help explain the contradictory results described in previous studies (14, 15, 17-24). Specifically, allergic asthma showed a small or insignificant effect on cancer of bronchi and lung in contrast to chronic obstructive asthma, which shows a pattern similar to the COPD emphysema subtype that is consistent with the known association between COPD and lung cancer (41-43).

The different localization patterns of neoplasms related to asthma and COPD that were identified here are consistent with mechanistic paradigms of the development of malignancy. Both asthma and COPD can affect the level of systemic inflammatory markers, hypoxia, and the composition and organization of the extracellular matrix (ECM) (44). Emerging data indicate that ECM has a crucial role in metastasis. Increased expression of genes encoding proteins that mediate ECM remodeling has been associated with increased mortality in patients with breast, lung, and gastric cancers (44-46). The results reported here that show an increased risk of breast, gastrointestinal, and lung cancer associated with asthma may be a result of increased expression of these proteins. The higher odds of malignant breast cancer and lower odds of lung and bronchial cancer with allergic asthma, compared to chronic bronchitis, bronchiectasis, and chronic airway obstruction, also support this hypothesis. Interestingly, the results show an association between allergic asthma and malignancies of ductal structures, including breast and the intrahepatic biliary system. Both elevated (47) and reduced risks (23) of lymphatic and hematopoietic cancer and asthma have been reported. Our results show increased odds of lymphatic and hematopoietic cancers in the presence of allergic asthma, suggesting an oncogenic relationship between asthma and lymphatic and hematopoietic cancer (48, 49).

The use of administrative data is constrained by the lack of clinical depth and potential coding errors. For example, acquiring accurate data on the onset of asthma and some confounder such as obesity was not possible with this dataset. Moreover, designing cohorts based on patients who had hospitalization encounters could have limited generalizability. In this study, using both primary and secondary diagnoses to identify asthma resulted in identifying asthma at a prevalence comparable to the general population indicating the methodology was sound. The use of ICD9 codes also limited the analysis of disease terminology that is not recent. Exposure to inhaled corticosteroid (ICS), and oral corticosteroid (OCS) therapy could confound the identified associations. However, corticosteroids have not been associated with thyroid or parathyroid neoplasms, suggesting it is unlikely that these medications are confounding our findings. Therefore, attributing the higher odds of neoplasia to corticosteroids needs further investigation.

This study provides the basis for future study design by generating common ORs from a large cohort that can be used in power calculations in future studies. As HCUP data does not contain outpatient medical office visits, our study was limited to inpatient and emergency department records, which usually contain records for patients with more severe asthma or later stages of COPD. Still, employing proper methods on these large datasets representative of the population at large provides the opportunity to improve the signal to noise ratio. In this study, the large cohort population allowed us to capture the relationship between allergic asthma and benign neoplasms to uncover the differences between obstructive and allergic asthma in the development of lung cancer.

In conclusion, these results show the value of using new methods of knowledge discovery, CICT, along with standard epidemiological methods to reveal signals that are invisible in the study of small datasets. We suggest that, in the era of data abundance, data-driven mass production, validation, our approach can be used to discover potential causal relationships in big clinical datasets. The uniformity of data preparation, analysis, and replication of 208 individual studies in our research makes the results readily comparable. This type of research could be used in parallel with review articles and meta and systematic analyses that integrate evidence from multiple sources. Our current analyses suggest a potential link between asthma, COPD, and neoplasms. Although the molecular mechanisms that underlie these links remain poorly understood and further research needs to be conducted, these studies suggest causal associations between asthma and neoplasms could be linked through biological processes.

**Appendix Table 1:** Pooled odds ratios for cancers in asthma and COPD by subtype. In exposure description, BN and MN correspondingly stand for ‘benign neoplasm’ and ‘malignant neoplasm’.

| ICD9CM Code | Exposure<br>Event | Asthma (493.*) | Asthma, unspecified (493.9) | Extrinsic asthma (493.0) | Chronic obstructive asthma (493.2) | Chronic bronchitis (491) | Bronchiectasis (494) | Chronic airway obstruction, not elsewhere classified (496) | Emphysema (492) |
| --- | --- | --- | --- | --- | --- | --- | --- | --- | --- |
| 210.2 | BN of major salivary glands | 1.4(1.26-1.56) | 1.29(1.14-1.45) | 2.73(1.78-4.2) | 2.04(1.71-2.43) | 1.93(1.68-2.23) | 2.11(1.31-3.4) | 2.14(1.93-2.38) | 2.66(2.1-3.37) |
| 227.3 | BN of pituitary gland and craniopharyngeal duct | 1.49(1.4-1.58) | 1.53(1.44-1.63) | 2.50(1.96-3.20) | 1.56(1.4-1.74) | 1.11(1.01-1.23) | 1.23(0.87-1.73) | 1.36(1.27-1.46) | 1.18(0.97-1.43) |
| 227.1 | BN of parathyroid gland | 1.44(1.33-1.55) | 1.41(1.3-1.53) | 2.43(1.77-3.35) | 1.75(1.54-2) | 1.25(1.11-1.41) | 1.77(1.23-2.55) | 1.69(1.56-1.83) | 1.92(1.58-2.33) |
| 225.2 | BN of cerebral meninges | 1.56(1.49-1.62) | 1.44(1.38-1.51) | 1.73(1.40-2.14) | 2.53(2.37-2.69) | 2.16(2.04-2.27) | 2.91(2.48-3.42) | 2.38(2.29-2.48) | 2.27(2.05-2.51) |
| 226 | BN of thyroid glands | 1.54(1.44-1.65) | 1.57(1.46-1.69) | 1.87(1.32-2.64) | 1.56(1.36-1.78) | 1.18(1.05-1.33) | 1.78(1.26-2.52) | 1.26(1.15-1.37) | 1.34(1.08-1.68) |
| 153.4 | MN of cecum | 1.17(1.09-1.24) | 0.99(0.92-1.07) | 1.33(0.95-1.85) | 2.15(1.96-2.36) | 1.86(1.72-2.01) | 2.82(2.26-3.53) | 2.79(2.66-2.94) | 3.18(2.83-3.57) |
| 153.6 | MN of ascending colon | 1.20(1.13-1.27) | 1.03(0.96-1.1) | 1.15(0.82-1.6) | 2.24(2.06-2.44) | 2.02(1.88-2.16) | 2.79(2.27-3.44) | 2.81(2.68-2.95) | 2.99(2.67-3.35) |
| 153.9 | MN of colon, unspecified site | 1.25(1.21-1.3) | 1.11(1.06-1.16) | 1.29(1.05-1.58) | 2.35(2.23-2.48) | 2.24(2.15-2.34) | 2.33(2.01-2.70) | 2.88(2.8-2.97) | 3.11(2.89-3.34) |
| 235.2 | Neoplasm of uncertain behavior of stomach, intestines, and rectum | 1.75(1.62-1.89) | 1.48(1.36-1.62) | 1.74(1.13-2.67) | 3.47(3.1-3.87) | 3.53(3.23-3.85) | 3.01(2.19-4.14) | 3.89(3.64-4.15) | 3.75(3.20-4.40) |
| 153.0 | MN of hepatic flexure | 1.31(1.17-1.47) | 1.12(0.98-1.28) | 1.67(0.95-2.95) | 2.51(2.13-2.97) | 2.23(1.94-2.56) | 3.19(2.13-4.77) | 3.29(3.01-3.6) | 3.77(3.06-4.63) |
| 153.1 | MN of transverse colon | 1.18(1.08-1.29) | 0.95(0.86-1.06) | 1.40(0.89-2.20) | 2.48(2.19-2.8) | 2.25(2.04-2.49) | 2.74(2.00-3.76) | 2.99(2.79-3.2) | 3.35(2.86-3.93) |
| 155.1 | MN of intrahepatic bile ducts | 1.00(0.89-1.11) | 0.92(0.82-1.05) | 1.75(1.09-2.82) | 1.62(1.36-1.93) | 1.21(1.03-1.41) | 2.07(1.35-3.18) | 1.87(1.7-2.07) | 2.26(1.8-2.83) |
| 156.1 | MN of extrahepatic bile ducts | 0.92(0.8-1.06) | 0.79(0.67-0.92) | 0.90(0.40-2.01) | 1.78(1.45-2.18) | 1.26(1.04-1.51) | 3.01(1.96-4.63) | 2.08(1.85-2.33) | 2.46(1.89-3.2) |
| 174.5 | MN of lower-outer quadrant of female breast | 1.58(1.39-1.79) | 1.57(1.37-1.8) | 2.23(1.23-4.04) | 1.89(1.51-2.38) | 1.46(1.2-1.79) | 0.58(0.19-1.80) | 1.6(1.38-1.85) | 2.04(1.46-2.86) |
| 174.4 | MN of upper-outer quadrant of female breast | 1.55(1.46-1.64) | 1.53(1.44-1.63) | 1.98(1.49-2.64) | 1.8(1.62-2.01) | 1.26(1.14-1.39) | 1.37(0.98-1.92) | 1.49(1.39-1.6) | 1.84(1.56-2.16) |
| 162.2 | MN of main bronchus | 4.64(4.29-5.01) | 2.65(2.4-2.93) | 2.32(1.37-3.92) | 13.73(12.57-14.99) | 22.31(20.91-23.81) | 10.75(8.44-13.7) | 21.5(20.3-22.77) | 33.78(30.96-36.86) |
| 162 | MN of trachea, bronchus, and lung | 3.08(3.02-3.14) | 1.87(1.82-1.91) | 1.76(1.55-2.00) | 8.69(8.5-8.89) | 13.95(13.73-14.18) | 9.2(8.69-9.74) | 14.87(14.67-15.07) | 25.35(24.79-25.92) |
| 162.4 | MN of middle lobe, bronchus or lung | 3.88(3.51-4.28) | 2.45(2.16-2.77) | 1.70(0.81-3.57) | 11.02(9.82-12.36) | 14.96(13.71-16.32) | 12.52(9.53-16.45) | 16.97(15.78-18.25) | 29.65(26.55-33.12) |
| 162.8 | MN of other parts of bronchus or lung | 3.36(3.18-3.55) | 2.09(1.94-2.24) | 1.36(0.88-2.11) | 9.42(8.82-10.05) | 15.55(14.85-16.28) | 9.99(8.5-11.75) | 15.72(15.11-16.35) | 25.01(23.49-26.64) |
| 188.9 | MN of bladder, part unspecified | 1.47(1.41-1.54) | 1.04(0.98-1.10) | 1.02(0.75-1.38) | 3.65(3.45-3.87) | 4.16(3.98-4.34) | 3.9(3.35-4.54) | 5.47(5.3-5.65) | 6.71(6.28-7.17) |
| 188.2 | MN of lateral wall of urinary bladder | 1.36(1.2-1.54) | 0.93(0.79-1.09) | 0.66(0.25-1.77) | 3.37(2.88-3.94) | 3.7(3.28-4.18) | 3.32(2.16-5.1) | 5.38(4.95-5.84) | 5.91(4.93-7.09) |
| 196.2 | Secondary and unspecified MN of intra-abdominal lymph nodes | 0.99(0.95-1.02) | 0.89(0.86-0.93) | 1.26(1.05-1.52) | 1.49(1.4-1.58) | 1.2(1.14-1.26) | 1.78(1.53-2.07) | 1.99(1.93-2.06) | 2.39(2.22-2.57) |
| 196 | Secondary and unspecified MN of lymph nodes | 1.28(1.25-1.30) | 1.13(1.11-1.16) | 1.54(1.39-1.71) | 2.02(1.96-2.09) | 2.06(2-2.11) | 2.2(2.02-2.39) | 2.73(2.68-2.78) | 3.85(3.72-4) |
| 198 | Secondary MN of other specified sites | 1.25(1.23-1.28) | 1.05(1.03-1.07) | 1.13(1.03-1.25) | 2.33(2.27-2.39) | 2.86(2.81-2.91) | 2.35(2.19-2.51) | 3.82(3.78-3.87) | 5.08(4.95-5.22) |
| 197 | Secondary MN of respiratory and digestive systems | 1.23(1.21-1.25) | 1.04(1.02-1.06) | 1.20(1.10-1.31) | 2.22(2.17-2.27) | 2.61(2.56-2.66) | 2.3(2.16-2.46) | 3.35(3.31-3.39) | 4.25(4.14-4.37) |
| 238.7 | Other lymphatic and hematopoietic tissues | 2.16(2.11-2.21) | 1.85(1.80-1.90) | 2.05(1.80-2.33) | 4.51(4.37-4.66) | 4.92(4.8-5.04) | 7.18(6.7-7.69) | 4.92(4.82-5.01) | 5.63(5.39-5.88) |

**Appendix table 2:** ICD9CM codes for cancers, asthma and COPD.

| ICD-9-CM | Description |
| --- | --- |
| 493.0 | Extrinsic asthma |
| 493.2 | Chronic obstructive asthma |
| 493.9 | Asthma, unspecified |
| 490 | Bronchitis, not specified as acute or chronic |
| 491 | Chronic bronchitis |
| 492 | Emphysema |
| 494 | Bronchiectasis |
| 496 | Chronic airway obstruction, not elsewhere classified |
| 210.2 | Benign neoplasm of major salivary glands |
| 227.3 | Benign neoplasm of pituitary gland and craniopharyngeal duct |
| 227.1 | Benign neoplasm of parathyroid gland |
| 225.2 | Benign neoplasm of cerebral meninges |
| 226 | Benign neoplasm of thyroid glands |
| 153.4 | Malignant neoplasm of cecum |
| 153.6 | Malignant neoplasm of ascending colon |
| 153.9 | Malignant neoplasm of colon, unspecified site |
| 235.2 | Neoplasm of uncertain behavior of stomach, intestines, and rectum |
| 153.0 | Malignant neoplasm of hepatic flexure |
| 153.1 | Malignant neoplasm of transverse colon |
| 174.5 | Malignant neoplasm of lower-outer quadrant of female breast |
| 174.4 | Malignant neoplasm of upper-outer quadrant of female breast |
| 155.1 | Malignant neoplasm of intrahepatic bile ducts |
| 156.1 | Malignant neoplasm of extrahepatic bile ducts |
| 162.2 | Malignant neoplasm of main bronchus |
| 162 | Malignant neoplasm of trachea, bronchus, and lung |
| 162.4 | Malignant neoplasm of middle lobe, bronchus or lung |
| 162.8 | Malignant neoplasm of other parts of bronchus or lung |
| 238.7 | Neoplasm of uncertain behavior of other lymphatic and hematopoietic tissues |
| 188.9 | Malignant neoplasm of bladder, part unspecified |
| 188.2 | Malignant neoplasm of lateral wall of urinary bladder |
| 180.8 | Malignant neoplasm of other specified sites of cervix |
| 196.2 | Secondary and unspecified malignant neoplasm of intra-abdominal lymph nodes |
| 196 | Secondary and unspecified malignant neoplasm of lymph nodes |
| 198 | Secondary malignant neoplasm of other specified sites |
| 197 | Secondary malignant neoplasm of respiratory and digestive systems |

## Appendix 3. Coarsened Exact Matching (CEM)

CEM applies an “exact matching” algorithm on coarsened variables and produces strata of matched case-control groups. We manually coarsened age into five categories: younger than 30, 30-49, 50-69, 70-89, and 90 and older. Also, the race was categorized as white, black, Hispanic, and other (including Asian or Pacific Islander, Native American, and Other in the original HCUP data). CEM generated weights were used in downstream analysis. Compared to other approximate matching methods, CEM shows superior performance in large datasets and comparable or better results with a controllable imbalance rate (34-36, 50) and includes the maximum possible number of controls in each stratum. CEM created 96 strata for combinations of accounted confounders in each cohort. For all cohorts, all patients in case groups had at least one matched person among controls. L1 statistics is a nonparametric measure that quantifies imbalance by comparing frequencies of the two groups across each of the strata. Values of L1 vary between 0 and 1, where values close to zero indicate perfect matching.

## Appendix 4. Advantages of CMH common odds ratio

The pooled odds ratio works on the whole cohort and ignores the fact that differences in the distribution of baseline covariates can have an uncontrolled effect on the measure of association. To address this problem, we used Cochran-Mantel-Haenszel (CMH) (37, 38) common odds ratio. CMH common odds ratio calculates odds ratio in each matched subgroups of a population and combines the odds ratios using a weight factor based on the size of subgroups. We calculated CMH common odds ratio overall balanced subgroups, or strata, returned by CEM algorithm. Effectively, CMH common odds ratio gives the estimate of odds ratios on a population that has been balanced on relevant confounders, based on the following formula:

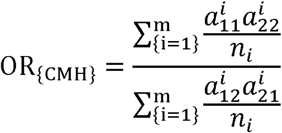

where *m* is the number of strata, 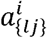 is the (*l, j*) entry of confusion matrix in *i*_*th*_ strata, and *n*_*i*_ is the number of patients in *i*_*th*_ stratum.

**Appendix Figure 2:**
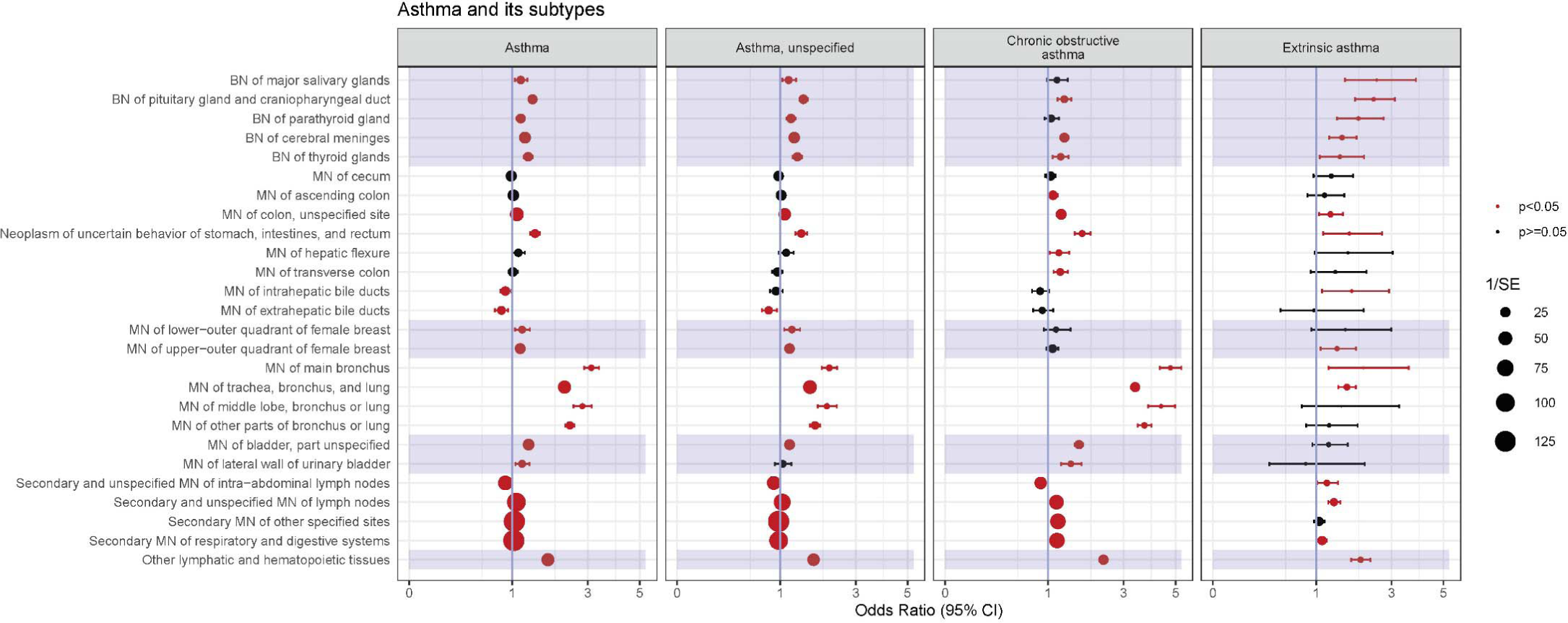
Results of individual asthma-neoplasm studies. Each line fragment shows the confidence interval for the odds ratio (dot) of a specific neoplasm(rows) with subtypes of asthma(columns). Significant OR that do not cross one (blue line) are in the red. Point sizes are inversely proportional to the standard error, so the larger points indicate more confident on the OR estimates and less standard error. In row labels, MN stands for malignant neoplasm, and BN stands for benign neoplasm.

**Appendix Figure 3:**
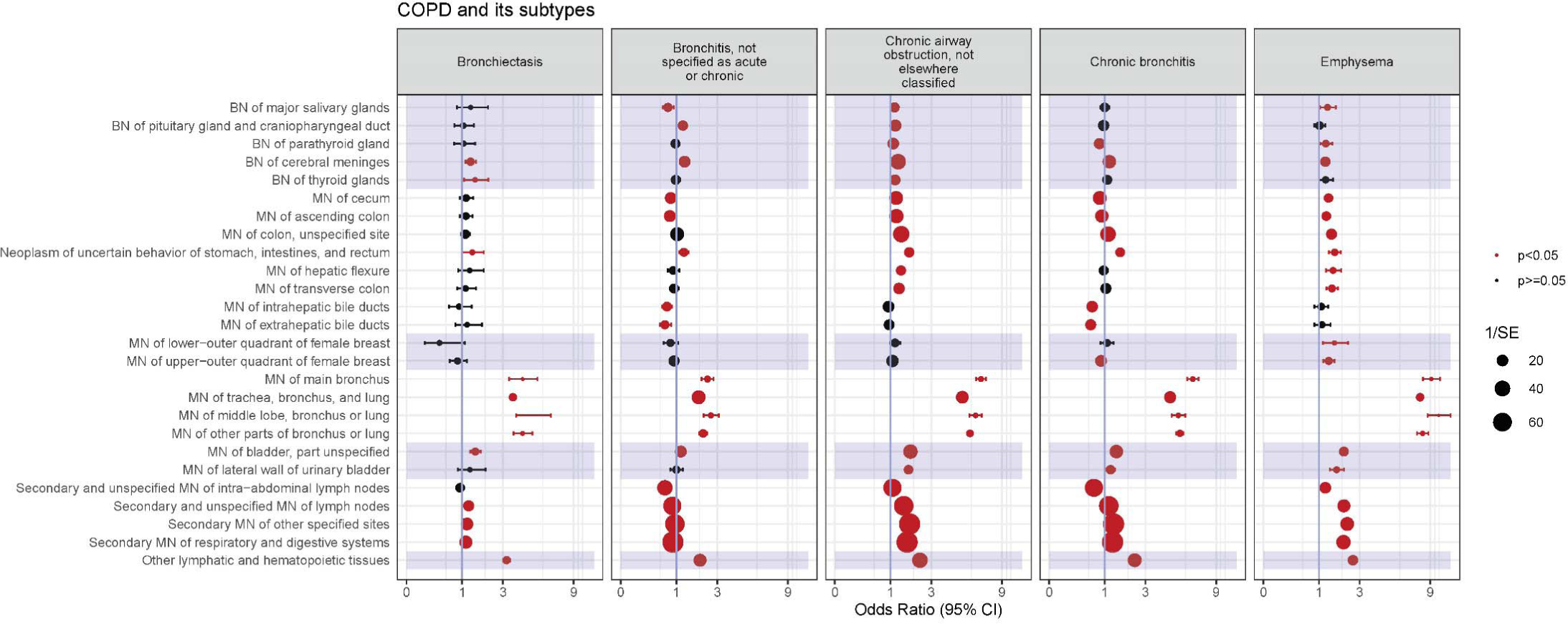
Results of individual COPD-neoplasm studies. Each line fragment shows the confidence interval for the odds ratio (black dot) of a specific neoplasm(rows) with subtypes of COPD(columns). Significant OR that do not cross one (blue line) are in the red. Point sizes are inversely proportional to the standard error, so the larger points indicate more confident on the OR estimates and less standard error. In row labels, MN stands for malignant neoplasm, and BN stands for benign neoplasm.

